# Loss of cBAF Complex Confers Sorafenib Resistance in Liver Cancer

**DOI:** 10.64898/2026.06.11.731725

**Authors:** Jacqueline A. Brinkman, Margarita Dzama-Karels, Max Bucklan, Carl J. Manner, Mallory Sokolowski, Jesse Raab

**Affiliations:** Lineberger Comprehensive Cancer Center, University of North Carolina, Chapel Hill, NC; Curriculum in Genetics and Molecular Biology, University of North Carolina, Chapel Hill, NC; Department of Genetics, University of North Carolina, Chapel Hill, NC

## Abstract

Sorafenib resistance limits the clinical benefit of first-line therapy in hepatocellular carcinoma (HCC), yet the epigenetic mechanisms underlying this resistance remain poorly understood. Using a chromatin-focused CRISPR/Cas9 dropout screen in HepG2 cells, we identified subunits of the canonical BAF (cBAF) complex — ARID1A, ARID1B, and SMARCC1 — as functional drivers of sorafenib resistance, while loss of the PBAF-specific subunit ARID2 had no effect, implicating cBAF specifically rather than SWI/SNF broadly. ARID1A and ARID1B mutations are associated with significantly worse overall survival in TCGA hepatocellular carcinoma cohorts. To define the underlying mechanism, we performed CUT&RUN profiling and RNA-seq in ARID1B-knockout HepG2 cells under sorafenib treatment. ARID1B loss triggered selective depletion of H3K4me3 and H3K27ac at 74 cBAF-dependent regulatory elements, 72% of which were directly occupied by ARID1B in wild-type cells. These sites were enriched for FOXA1 and HNF4α motifs, consistent with disruption of hepatocyte lineage regulatory chromatin. Direct profiling in HLF hepatocellular carcinoma cells confirmed that 179 FOXA1 binding sites were lost specifically under the combined perturbation of ARID1B loss and sorafenib treatment, with neither condition alone sufficient to drive this effect. Transcriptionally, ARID1B loss induced epithelial-mesenchymal transition programs and suppressed MYC targets, E2F targets, and mTORC1 signaling. These findings define a cBAF–FOXA1 regulatory axis that maintains hepatocyte lineage identity under kinase inhibitor stress, and whose disruption drives epigenetic reprogramming toward a mesenchymal drug-tolerant state.

**Graphical Abstract:** 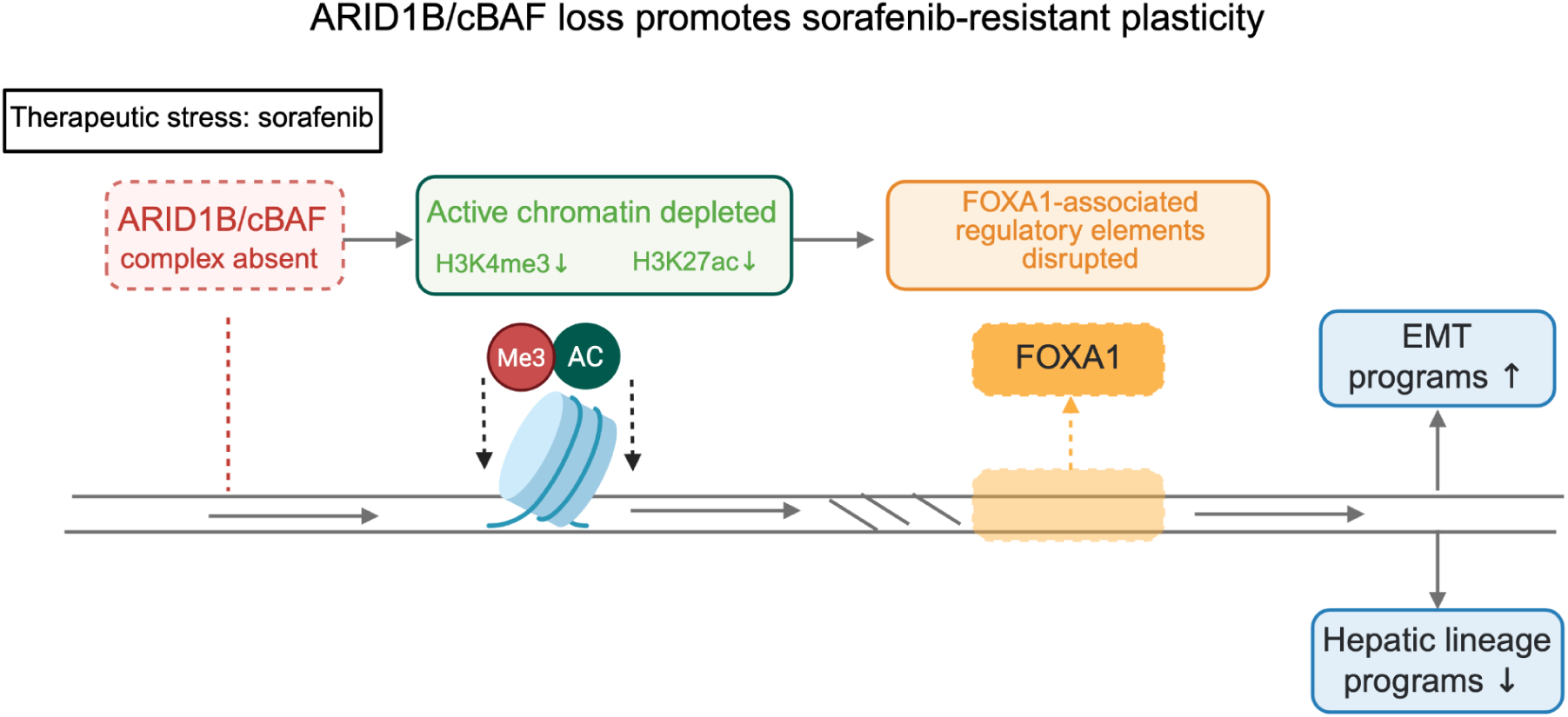

## Introduction

Chromatin remodeling complexes are among the most frequently altered epigenetic regulators in human cancer, with mutations in SWI/SNF complex subunits occurring in approximately 20% of all tumors^1,2^. The SWI/SNF family encompasses three major assemblies, canonical (cBAF), polybromo-associated (PBAF), and non-canonical (ncBAF)^3^, each defined by distinct subunit composition and performing non-redundant functions in regulating chromatin accessibility ^4^, enhancer activity^5–7^, and transcription factor occupancy at lineage-specific regulatory elements ^8,9^. Individual SWI/SNF subunits can function as tumor suppressors ^10–12^ or context-dependent oncogenes depending on cancer type ^13^, and their disruption can alter not only transcriptional regulation but also cellular responses to therapeutic stress ^14,15^. Ongoing clinical trials are evaluating degraders targeting SMARCA2, dual SMARCA2/SMARCA4, and BRD9; ^16^ however, how loss of individual SWI/SNF complex components shapes sensitivity to existing therapies remains poorly understood. This represents a critical gap in translating epigenetic biology into patient benefit.

Hepatocellular carcinoma (HCC) is the most common primary liver malignancy and the third leading cause of cancer-related mortality worldwide.^17^ Sorafenib, a multikinase inhibitor targeting the RAF-MEK-ERK and VEGFR signaling, was the first approved first-line therapy for advanced HCC.^18^ However, sorafenib only provides a modest survival benefit and resistance is nearly universal.^19^ The molecular basis of sorafenib resistance is heterogeneous and incompletely understood, with proposed mechanisms including activation of bypass signaling pathways, epithelial-mesenchymal transition (EMT), and metabolic reprogramming.^20–22^ Although multiple genetic and functional screens have identified resistance regulators, epigenetic mechanisms are comparatively underexplored ^21,23–26^. The high frequency of chromatin regulator mutations in HCC suggest that a dysregulated epigenome could contribute to resistance. ARID1A and ARID1B, mutually exclusive DNA-targeting subunits of the cBAF complex, are among the most recurrently altered genes, collectively mutated in 15–20% of cases ^27,28^ .

Pioneer transcription factors including FOXA1, FOXA2, and HNF4α establish and maintain the hepatocyte-specific regulatory landscape by binding nucleosomal DNA and licensing downstream transcription factor occupancy.^29,30^ cBAF complexes cooperate with lineage-determining transcription factors to maintain nucleosome-depleted regulatory regions and support enhancer activity, and disruption of this cooperation has been linked to loss of lineage identity in multiple contexts ^31,32^. Whether cBAF integrity is required to preserve pioneer factor occupancy during therapeutic stress, and whether cBAF loss can reprogram the hepatocyte regulatory landscape toward a drug-tolerant state, remains an open question.

Here we show that loss of cBAF subunits ARID1A, ARID1B, and SMARCC1 confers sorafenib resistance in HCC. Using a chromatin-focused CRISPR/Cas9 screen followed by chromatin profiling and transcriptomic analysis across HCC cell line models, we identify ARID1B as a regulator of sorafenib response and define an epigenetic mechanism centered on disruption of the FOXA pioneer factor program. ARID1B loss under sorafenib treatment triggered selective collapse of active chromatin at cBAF-dependent hepatocyte lineage regulatory elements, reduced FOXA1 occupancy at a discrete set of binding sites, and transcriptional reprogramming toward an EMT-associated drug-tolerant state. These findings identify the cBAF–FOXA1 regulatory axis as a determinant of sorafenib sensitivity and suggest that maintenance of pioneer factor-bound chromatin represents a key vulnerability in cBAF-deficient HCC.

## Materials and Methods

### Cell culture and drug treatment

Human hepatocellular carcinoma cell lines HLF (CVCL_2947) and HepG2 (CVCL_0027), and Lenti-X 293 cells (CVCL_0045), were obtained from ATCC. Cell line identity was authenticated by ATCC at the time of purchase. Cells were maintained at 37°C in a humidified incubator with 5% CO₂ in Dulbecco’s modified Eagle medium (DMEM) supplemented with 10% fetal bovine serum and 1% penicillin–streptomycin (Gibco). Cells were passaged every 3–4 days and replated in fresh medium.

## Drug treatment and cell viability assays

Trypan blue-excluding viable cells were seeded in 96-well plates at 10,000 cells per well in 150 μl complete medium. For IC₅₀ determination, sorafenib and donafenib (Selleckchem, S1040 and S9621) were prepared by two-fold serial dilution across nine concentrations, with 50 μM as the highest concentration. Vehicle-treated cells received DMSO. Cells were treated for 72 h, with technical and biological triplicates performed for each condition.

Cell viability was measured using the CellTiter-Glo Luminescent Cell Viability Assay (Promega) according to the manufacturer’s instructions. Briefly, CellTiter-Glo reagent was added directly to each well at a 1:1 ratio with culture medium and incubated for 10 min at room temperature with gentle agitation. Luminescence was measured using a Cytation 5 plate reader (BioTek). Viability values were normalized to the corresponding DMSO-treated control.

### Immunoblotting

Cells treated with sorafenib (Selleckchem, S1040) or vehicle control (DMSO) were lysed in RIPA buffer at a ratio of 1 × 10⁶ cells per 30 μl lysis buffer. RIPA buffer contained 150 mM NaCl, 1.0% IGEPAL CA-630, 0.5% sodium deoxycholate, 0.1% SDS and 50 mM Tris, pH 8.0. Equal amounts of protein were loaded onto 4–15% Mini-PROTEAN TGX precast gels (Bio-Rad, #456-1034), separated by SDS–PAGE and transferred to PVDF membranes using a Trans-Blot Turbo transfer system (Bio-Rad) according to the manufacturer’s protocol.

Membranes were blocked for 1 h at room temperature in a 1:1 mixture of Intercept blocking buffer (LI-COR) and TBS. TBS contained 200 mM Tris and 1500 mM NaCl. Membranes were cut as needed to allow independent antibody incubations and were incubated overnight at 4°C with gentle agitation in primary antibodies diluted in antibody buffer. The following primary antibodies were used: anti-β-actin (1:5000, Cell Signaling Technology, #3700) and anti-ARID1B (1:1000, Cell Signaling Technology, #65747, clone ID: E1U7D).

Membranes were washed three times for 10 min each at room temperature in TBST containing 0.1% Tween-20, then incubated for 1 h at room temperature with fluorescent secondary antibodies diluted 1:20,000 in antibody buffer. The following secondary antibodies were used: goat anti-rabbit 800 (Amersham, #PA45004) and goat anti-mouse 680 (Amersham, #PA45002). Membranes were imaged using an Odyssey CLx imaging system (LI-COR), and immunoblot signal was quantified using Image Studio software (LI-COR).

### CRISPR/Cas9 screen

#### CRISPR/Cas9 library design and cloning

A focused CRISPR/Cas9 sgRNA library was designed using GUIDES (http://guides.sanjanalab.org/) and a custom-curated set of chromatin regulatory genes (Table S1). The library contained approximately 6000 sgRNAs, including ∼800 non-targeting controls, targeting approximately 750 transcription factors and chromatin regulatory genes, including components of SWI/SNF and PRC1/2 complexes, as well as SIRT, HDAC, DNMT and MBD family members. Oligonucleotides were synthesized by Twist Bioscience and cloned into pLentiCRISPRv2 (Addgene, #52961) as previously described^33^.

#### Lentivirus production and transduction

Lentivirus containing the pooled CRISPR/Cas9 sgRNA library was generated by transfecting HEK293T cells with psPAX2, pLentiCRISPRv2-sgRNA library and pMD2.G at a 2:2:1 ratio. Viral supernatant was harvested 72 h after transfection.

Cells were transduced at a multiplicity of infection of 0.25–0.3 to favor single viral integration events and selected with 1 μg/ml puromycin for 7 days to remove uninfected cells.

#### Pooled screen and sequencing library preparation

Baseline cell pellets containing at least 5 × 10⁶ cells were harvested after puromycin selection for genomic DNA extraction using the Monarch Genomic DNA Purification Kit (NEB #T3010). Following selection, 1.5 × 10⁶–3 × 10⁶ cells were split between vehicle control (DMSO) and two sorafenib concentrations (2.5 μM and 4 μM; the higher concentration approximates the 96 h IC50 in HepG2 cells) then maintained under selective pressure for 28 days, corresponding to greater than 500-fold library coverage. Drug-containing medium was refreshed every 3 days, and cells were passaged as needed based on density while maintaining library representation.

After 28 days, genomic DNA was extracted as described above. High-throughput sequencing libraries were generated by PCR amplification of integrated sgRNA sequences using primers that added Illumina adapters and sample barcodes. Libraries were sequenced as single-end 75-bp reads on an Illumina NextSeq 500.

#### CRISPR/Cas9 screen analysis

CRISPR/Cas9 screen data were analyzed using the MAGeCK^34^ pipeline, with non-targeting sgRNAs used as controls. Gene-level enrichment and depletion were assessed using the MAGeCK robust ranking algorithm for each contrast. Robust ranking algorithm scores were compared across treatment conditions in R. For individual sgRNA-level analyses, read counts were normalized for sequencing depth before visualization.

#### RNA-seq

Cells were harvested in biological replicates by pelleting 1 × 10⁶ cells for 1 min at 500 × g at room temperature. Total RNA was isolated using the Monarch Total RNA Miniprep Kit (NEB #T1010) with on-column DNase I digestion.

#### RNA-seq library preparation

mRNA-seq libraries were prepared from 750 ng total RNA using the KAPA mRNA HyperPrep Kit (Roche, #08098123702) according to the manufacturer’s instructions. RNA was fragmented for 6 min at 94°C to generate libraries with an expected insert size of 200–300 bp. Adapter ligation was performed using 7 μM adapter stock, and libraries were amplified for 9 PCR cycles. Libraries were quantified, pooled, and sequenced as single-end 75-bp reads on an Illumina NextSeq 100.

#### RNA-seq analysis

Raw sequencing reads were quantified against the human reference transcriptome GRCh38, Ensembl release 109, using Salmon^35^ with default parameters. Transcript-level quantifications were imported into R and summarized to gene-level counts using tximeta^36^ and tximport. Differential expression analysis was performed using DESeq2^37^ v1.46.0 with a full interaction model: genotype + treatment + genotype. Genes with fewer than 10 counts in at least three samples were excluded prior to modeling. Log₂ fold-change estimates were regularized using apeglm^38^ shrinkage.

The primary contrast of interest was the genotype-by-treatment interaction term: WT_DMSO, ARID1B-KO_DMSO, WT_sorafenib and ARID1B-KO_sorafenib, n = 3 biological replicates per condition, which identified genotype-specific transcriptional responses to sorafenib treatment. For ARID1B-knockout versus wild-type comparisons under sorafenib treatment, differentially expressed genes were defined using an adjusted P value threshold of 0.1. Gene set enrichment analysis was performed using fgsea^39^ and the MSigDB Hallmark gene set collection v2023.1. For overrepresentation analysis, significantly upregulated and downregulated genes were tested separately using fgsea::fora, with all expressed genes used as the background universe.

### CUT&RUN

#### CUT&RUN sample preparation

CUT&RUN was adapted from the EpiCypher protocol. For each sample, 200,000–500,000 cells were pelleted for 3 min at 300 × g at room temperature. Cells were washed twice in wash buffer containing 20 mM HEPES, pH 7.6, 150 mM NaCl, 0.5 mM spermidine and protease inhibitor cocktail. For each sample, 10 μl Concanavalin A beads (Bangs Laboratories, BP531) were washed twice in bead activation buffer containing 20 mM HEPES, pH 7.9, 10 mM KCl, 1 mM CaCl₂ and 1 mM MnCl₂. Cells were incubated with activated Concanavalin A beads in wash buffer for 5–10 min at room temperature.

Supernatant was removed, and bead-bound cells were resuspended in 50 μl wash buffer containing 0.05% digitonin and 2 mM EDTA. Primary antibodies were added, and samples were incubated overnight at 4°C. The following antibodies were used: anti-H3K4me3 (Cell Signaling Technology, #3076423), anti-H3K27ac (Cell Signaling Technology, #2561016) and rabbit IgG control (Cell Signaling Technology, #550038). Bead-bound cells were washed twice on a magnet using wash buffer containing 0.05% digitonin, hereafter referred to as digitonin wash buffer, and incubated in 50 μl digitonin wash buffer containing guinea pig anti-rabbit secondary antibody (1:100; Novus Biologicals, NBP1-72763) for 1 h at 4°C.

Bead-bound cells were washed as described above and incubated with 700 ng/ml pAG-MNase, purified in-house as previously described^40^, for 1 h at 4°C. Samples were washed four times with cold digitonin wash buffer and resuspended in 50 μl cold digitonin wash buffer on ice. MNase digestion was initiated by adding 1 μl of 100 mM CaCl₂, and samples were incubated on ice at 4°C for 30 min. Reactions were quenched by adding stop buffer containing 340 mM NaCl, 20 mM EDTA, 4 mM EGTA, 0.05 μg/μl RNase A (Thermo Fisher Scientific, EN0531) and 0.1% Triton X-100. Samples were incubated at 37°C for 30 min in a thermocycler, and DNA was purified using the Zymo DNA Clean & Concentrator-25 Kit (Zymo Research, D4014).

#### CUT&RUN library preparation

CUT&RUN libraries were prepared using the KAPA HyperPrep Kit (Roche, #07962363001) with modifications. End repair and A-tailing reactions were performed at 12°C for 15 min, 37°C for 15 min and 58°C for 45 min. Adapters were ligated for 1 h using 5 μl of 750 nM Roche dual-index adapter stock (Roche, #08861919702). Libraries were cleaned twice using 1.1× volumes of KAPA Pure Beads and amplified using KAPA HiFi PCR mix with the following cycling conditions: 98°C for 30 s; 14 cycles of 98°C for 15 s and 60°C for 10 s; followed by final cleanup using 1.2× volumes of KAPA Pure Beads (Roche, #07983298001). Libraries were quantified, pooled and sequenced as paired-end 50-bp or paired-end 150-bp reads on an Illumina NextSeq 1000 or NovaSeq X Plus 10B.

#### CUT&RUN read processing and peak calling

Read alignment and peak calling were performed using the laboratory’s Nextflow CUT&RUN pipeline^41,42^. Reads were aligned to the GRCh38 human reference genome using Bowtie2^43^. Peaks were called using MACS2 in paired-end mode with -f BAMPE. Consensus peaks across biological replicates were identified using MSPC^44^ with Fisher’s method and the following parameters: -r bio -w 1e-4 -s 1e-5 -c 2. Normalization factors were computed by the pipeline using csaw^45^ and retained for each sample.

#### Differential occupancy analysis

Read counts within merged consensus peak sets were quantified using Rsubread::featureCounts in paired-end mode. Peaks overlapping the hg38 CUT&RUN exclusion list were removed prior to analysis. Differential occupancy was assessed using DESeq2 with a ∼ condition design. csaw normalization factors generated by the pipeline were applied as DESeq2 size factors in place of library-size normalization. Peaks with fewer than five counts in at least two samples were excluded prior to modeling. Peaks with adjusted P < 0.5 and absolute log₂ fold change > 0.5 were considered differentially occupied. The primary contrast was ARID1B-knockout versus wild-type cells under sorafenib treatment. Shared regions losing both H3K4me3 and H3K27ac were identified by overlapping significantly decreased H3K4me3 and H3K27ac peak sets using GenomicRanges::findOverlaps.

#### Motif enrichment analysis

*De novo* and known motif enrichment analyses were performed on differential peak sets using HOMER findMotifsGenome.pl v5.1 against the hg38 genome. Motif enrichment was performed using a 200-bp window centered on each peak, with repeat masking enabled using -size 200 -mask. Known motif enrichment was assessed using the HOMER vertebrate motif database.

#### Signal visualization

Condition-averaged BigWig files were generated from csaw scaled coverage tracks using deepTools^46^ bigwigAverage v3.5.6. Signal matrices were computed using deepTools computeMatrix reference-point with a ±3-kb window centered on peak midpoints and the following parameters: --referencePoint center --beforeRegionStartLength 3000 --afterRegionStartLength 3000 --binSize 50 --missingDataAsZero. Peaks overlapping the hg38 CUT&RUN exclusion list were excluded during matrix computation using --blackListFileName.

CUT&RUN signal tracks were visualized using the plotgardener R package (v1.12.0). Csaw scaled bigWig files were imported and displayed for a representative replicate at the KLF12 locus (chr13:74,081,000–74,085,000, hg38). Signal range was matched across all four conditions (0–32 reads per genomic content) to enable direct comparison.

#### Peak-to-gene linking

Differential CUT&RUN peaks were linked to nearby genes by extending peak coordinates 50 kb in each direction and identifying overlaps with gene bodies using GenomicRanges::findOverlaps. Gene annotations were obtained from TxDb.Hsapiens.UCSC.hg38.knownGene. Gene identifiers were mapped to Ensembl IDs and gene symbols using org.Hs.eg.db. Peaks were linked to the nearest expressed gene with a transcription start site within 50 kb of the peak center. This proximity-based strategy was used to nominate putative target genes for expression-level comparison and was not interpreted as definitive enhancer-gene assignment. Linked genes were intersected with RNA-seq differential expression results to identify genes with concordant chromatin and transcriptional changes. As a sensitivity analysis, we also evaluated links within 100 kb and observed similar qualitative trends.

#### FOXA1/2 occupancy analysis

CUT&RUN data for FOXA1 and FOXA2 in HLF cells were processed using the same alignment, peak calling and differential occupancy analysis framework described above for histone marks. The primary contrast was ARID1B-knockout versus wild-type cells under sorafenib treatment. Overlap between FOXA1/2 binding sites and shared regions losing both H3K4me3 and H3K27ac was assessed using GenomicRanges::findOverlaps after extending shared peak coordinates by 5 kb.

Genes near lost FOXA1 sites were identified using the 50-kb peak-to-gene linking strategy described above. Expression changes among genes linked to lost FOXA1 sites were compared with a random background set of 1000 genes not linked to lost FOXA1 sites using a two-sided Wilcoxon rank-sum test.

#### Data visualization and software

Box plots, volcano plots and scatter plots were generated in R using ggplot2^47^ v4.0.1. Gene labels were added using ggrepel^48^ v0.9.6 to minimize label overlap. Genome browser tracks were generated using plotgardener^49^ and display mean-normalized signal across biological replicates. All analyses were performed in R v4.4.0. Package versions were tracked using renv v1.2.1. Analysis notebooks are available at https://github.com/jraab/2026_sorafenib_resistance. Key package versions included DESeq2 v1.46.0, tximeta v1.24.0, Rsubread v2.20.0, GenomicRanges v1.58.0, fgsea v1.32.4, ggplot2 v4.0.2, deepTools v3.5.6 and HOMER v5.1.

## RESULTS

To define chromatin regulators that influence sorafenib response in liver cancer, we performed a focused CRISPR/Cas9 knockout screen targeting 737 chromatin regulatory genes in HepG2 cells. Cells were transduced with a lentiviral sgRNA library, selected with puromycin to remove uninfected cells, and harvested at baseline to capture the initial sgRNA distribution. Following selection, cells were assigned to vehicle or sorafenib treatment arms and cultured for 28 days in DMSO, 2.5 μM sorafenib, or 4 μM sorafenib, with 4 μM approximating the 96-hour IC₅₀ in HepG2 cells. Genomic DNA was then isolated, and sgRNA representation was quantified by sequencing to identify genes whose loss altered cellular fitness under sorafenib treatment (**Figure 1A**).

**Figure 1:**
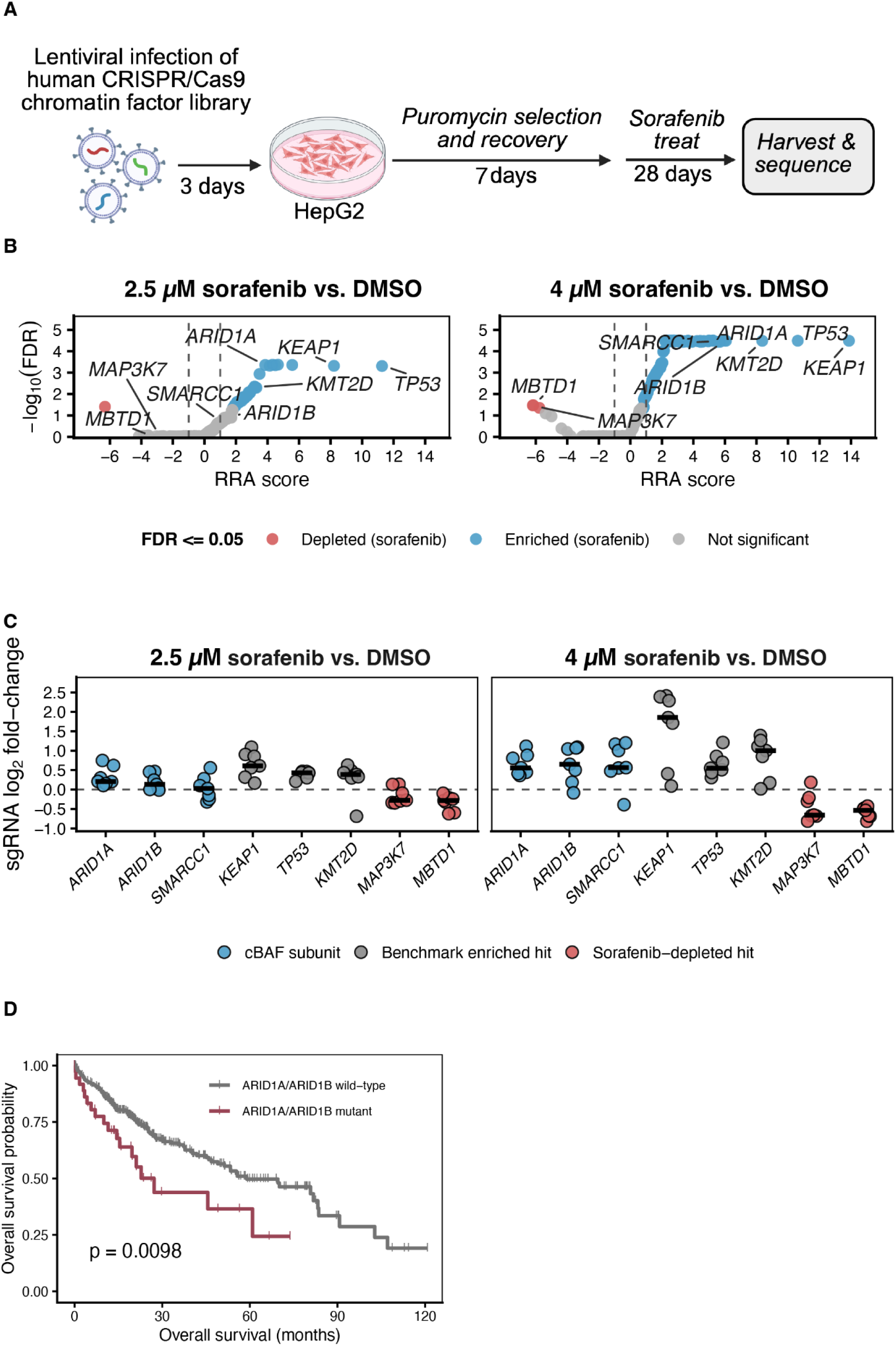
A chromatin-focused CRISPR screen identifies cBAF as a regulator of sorafenib resistance. **(A)** Schematic of the chromatin-focused CRISPR/Cas9 dropout screen in HepG2 cells. Cells were transduced with a lentiviral sgRNA library targeting chromatin regulators, selected with puromycin and cultured under DMSO or sorafenib treatment. Genomic DNA was collected at baseline and after treatment for sgRNA sequencing. **(B)** Gene-level MAGeCK robust ranking algorithm analysis comparing HepG2 cells treated with 2.5 uM and 4 uM sorafenib versus DMSO. Each point represents one gene. The x-axis shows the RRA score, with positive values indicating enrichment under sorafenib selection and negative values indicating depletion. The y-axis shows −log₁₀(FDR). Genes with FDR ≤ 0.05 are colored blue if enriched under sorafenib and red if depleted under sorafenib; non-significant genes are shown in grey. Labeled genes include cBAF complex members and selected cancer-associated hits. **(C)** Individual sgRNA-level log₂ fold-change for selected screen hits in 2.5 uM and 4 uM sorafenib versus DMSO comparisons. Each point represents one sgRNA targeting the indicated gene. Crossbars indicate the median sgRNA log₂ fold-change per gene. Positive values indicate enrichment under sorafenib selection, whereas negative values indicate depletion. **(D)** Kaplan–Meier overall survival analysis of TCGA hepatocellular carcinoma patients stratified by ARID1A or ARID1B mutation status. Patients with tumors harboring mutations in either gene exhibited significantly worse overall survival than patients lacking mutations in both genes (log-rank p = 0.0098).

MAGeCK robust ranking algorithm analysis identified genes whose sgRNAs were selectively enriched or depleted under sorafenib treatment relative to DMSO. Across both sorafenib doses, sgRNAs targeting canonical cancer-associated genes, including KEAP1, TP53, and KMT2D, were enriched in sorafenib-treated populations, consistent with selective outgrowth of resistant cells. Components of the canonical BAF complex were also recovered among sorafenib-enriched hits, including ARID1A, ARID1B, and SMARCC1, with stronger enrichment observed at the 4 μM dose (**Figure 1B**). Individual sgRNA-level analysis confirmed concordant enrichment of guides targeting cBAF subunits under sorafenib selection (**Figure 1C**). These results suggest that loss of cBAF complex activity provides a selective advantage during sorafenib exposure and nominate ARID1B as a candidate regulator of sorafenib sensitivity.

To determine whether sorafenib-associated enrichment was specific to cBAF or reflected broader disruption of BAF-family chromatin remodelers, we next examined individual sgRNAs targeting PBAF-associated subunits. In contrast to sgRNAs targeting ARID1A, ARID1B, and SMARCC1, sgRNAs targeting the PBAF-specific subunits ARID2, PBRM1, and BRD7 were not enriched under sorafenib selection. These results support a cBAF-specific role in modulating sorafenib response (**Supplementary Figure S1A**).

Because ARID1A and ARID1B are recurrently mutated in hepatocellular carcinoma, we next asked whether these alterations were associated with clinical outcome. Using cBioPortal data from The Cancer Genome Atlas PanCancer hepatocellular carcinoma cohort, we found that tumors harboring ARID1A or ARID1B mutations were associated with significantly worse overall survival than tumors lacking mutations in both genes (median overall survival, 45.6 versus 80.7 months; log-rank P = 0.0098; **Figure 1D**). These findings identify cBAF loss as a candidate chromatin-based mechanism of sorafenib resistance and support further investigation of ARID1B-dependent regulation of drug response in hepatocellular carcinoma.

### ARID1B loss confers resistance to sorafenib and donafenib in liver cancer models

To validate the CRISPR screen findings, we generated ARID1B-knockout HepG2 cells using individual sgRNAs and confirmed depletion of ARID1B protein by immunoblotting (**Figure 2A**). ARID1B-knockout cells showed reduced sensitivity to sorafenib compared with control cells, reflected by a rightward shift in the dose–response curve (**Figure 2B**). A similar reduction in drug sensitivity was observed following treatment with donafenib, a structurally related multikinase inhibitor, indicating that the effect of ARID1B loss is not limited to sorafenib alone (**Figure 2C**).

**Figure 2:**
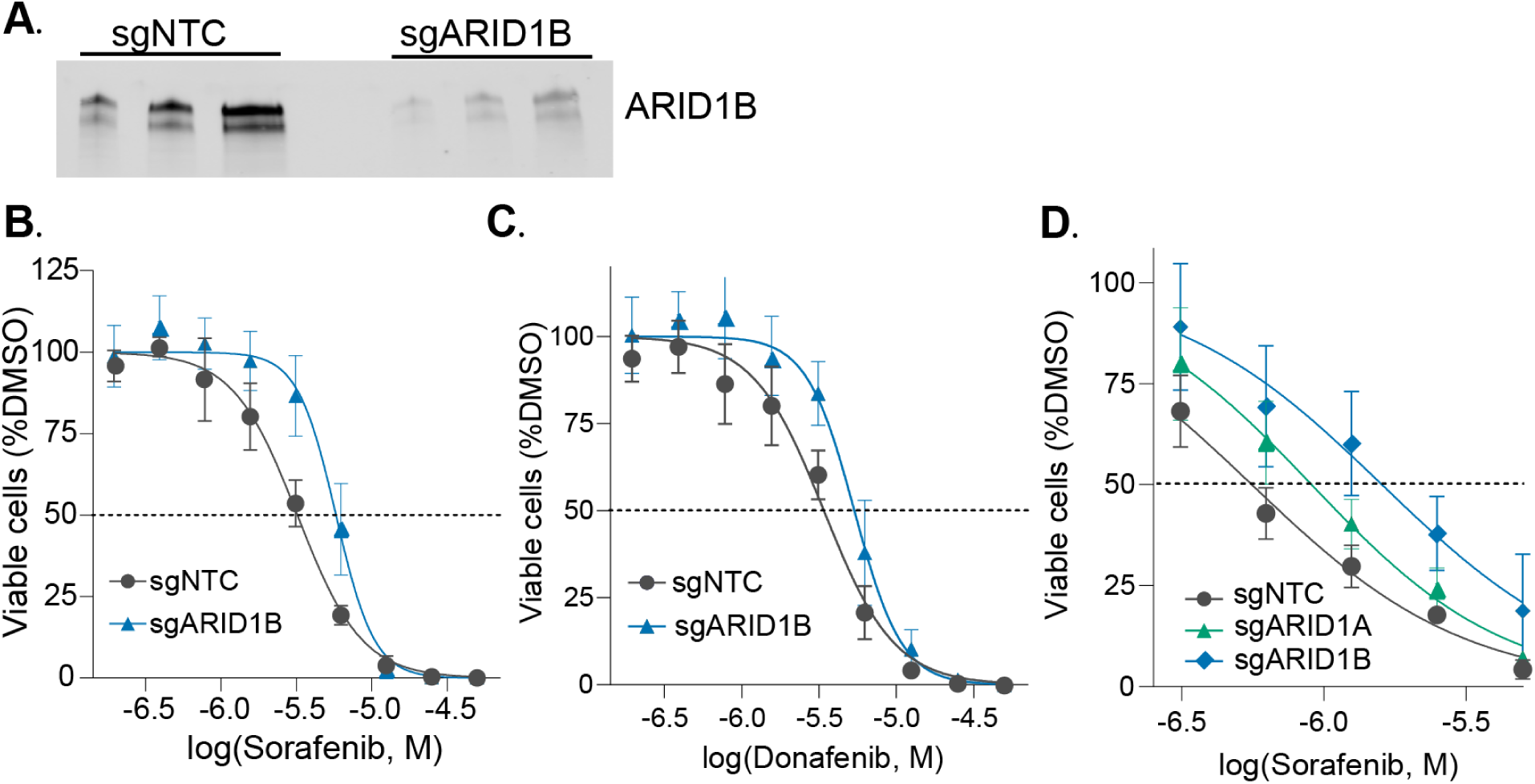
ARID1B loss confers sorafenib resistance across liver cancer models. **(A)** Generation and validation of ARID1B-knockout HepG2 cells. Immunoblot showing loss of ARID1B protein in knockout clones relative to wild-type controls. **(B)** Cell viability of wild-type/control and ARID1B-knockout HepG2 cells across a nine-point, two-fold serial dilution series (50 uM to 0.2 uM) for 72h of sorafenib. Viability was normalized to DMSO control. Data shown as mean ± SEM (n = 2 biological replicates, 3 technical replicates). **(C)** Cell viability of wild-type/control and ARID1B-knockout HepG2 cells treated with donafenib under identical conditions. **(D)** Cell viability of HLF control, ARID1A-knockout and ARID1B-knockout cells treated with increasing concentrations of sorafenib. Data are normalized to DMSO-treated controls and shown as mean ± SEM.

To determine whether this resistance phenotype was conserved in an additional liver cancer model, we generated ARID1A- or ARID1B-knockout HLF hepatocellular carcinoma cells and assessed their response to sorafenib. Knockout of either ARID1A or ARID1B increased cell viability under sorafenib treatment relative to control cells, supporting a conserved role for cBAF complex disruption in reducing sorafenib sensitivity across liver cancer models (**Figure 2D**).

### ARID1B loss remodels active chromatin landscapes during sorafenib treatment

To define how ARID1B loss alters chromatin regulation during the sorafenib response, we performed CUT&RUN profiling for H3K4me3 and H3K27ac in wild-type and ARID1B-knockout HepG2 cells treated with DMSO or 4 uM sorafenib for 72 hours (**Figure 3A**). Principal component analysis (PCA) demonstrated clear separation of samples by genotype and treatment condition, supporting the reproducibility of the dataset (**Supplemental Figure S2A-B**).

**Figure 3:**
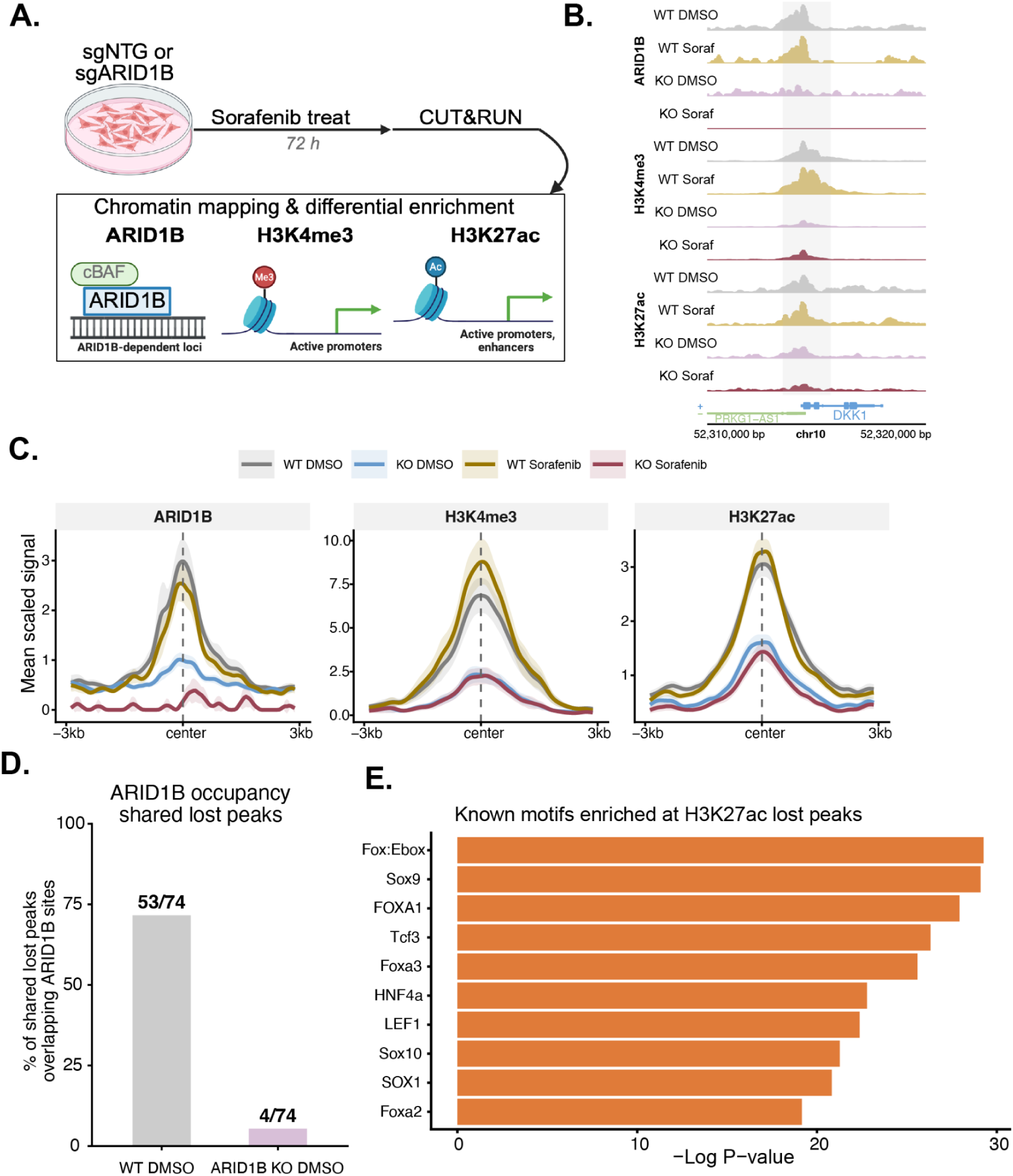
ARID1B loss depletes active chromatin at hepatocyte lineage regulatory elements during sorafenib treatment. **(A)** Schematic of CUT&RUN experimental design. Wild-type and ARID1B-knockout HepG2 cells were treated with DMSO or sorafenib and profiled for ARID1B, H3K4me3 and H3K27ac by CUT&RUN (n = 2 biological replicates per condition). **(B)** Representative genome browser tracks at the DKK1 locus showing normalized CUT&RUN signal for ARID1B, H3K4me3 and H3K27ac across the indicated conditions. Shaded region denotes a shared lost peak. **(C)** Profile plots showing scaled CUT&RUN signal for ARID1B, H3K4me3 and H3K27ac centered on 74 genomic regions that lose both H3K4me3 and H3K27ac in ARID1B-knockout cells under sorafenib treatment (±3 kb). Shading indicates ±1 SEM across peaks. **(D)** Percentage of shared lost peaks (n = 74) overlapping ARID1B CUT&RUN peaks in wild-type (53/74, 72%) and ARID1B-knockout (4/74, 5%) HepG2 cells. **(E)** Known motif enrichment analysis at H3K27ac using HOMER. Top 10 motifs ranked by −log10(p-value).

Differential peak analysis identified 336 H3K4me3 peaks and 557 H3K27ac peaks significantly decreased relative to wild-type cells (adjusted P < 0.1, |log2FC| > 0.5; **Supplemental Figure S2C-D**). The DKK1 locus illustrates this effect, showing selective loss of ARID1B, H3K4me3 and H3K27ac signal in knockout cells under sorafenib treatment (**Figure 3B**). Genomic annotation of differential peaks showed that H3K4me3 loss occurred predominantly at promoters (76%), whereas H3K27ac loss was enriched at intronic and distal intergenic regions (62%), consistent with disruption of distal enhancer-associated chromatin (**Supplemental Figure S2E**).

To identify high-confidence ARID1B-dependent regulatory elements, we intersected regions losing H3K4me3 with those losing H3K27ac and identified 74 shared sites. Profile plots centered on these shared lost regions showed marked depletion of H3K4me3 and H3K27ac signals specifically in ARID1B-knockout cells under sorafenib treatment, while wild-type cells retained stable or increased signal (**Figure 3C**). To test whether these sites were directly ARID1B-associated, we intersected the shared lost regions with ARID1B CUT&RUN peaks. 53 out of 74 shared lost peaks (72%) overlapped ARID1B binding sites in wild-type cells, compared with only 4 of 74 sites (5%) in ARID1B-knockout cells (**Figure 3D**), supporting the conclusion that these regions represent ARID1B-dependent regulatory elements.

Motif enrichment analysis at H3K27ac lost peaks revealed strong enrichment for Forkhead family motifs, including Fox:Ebox and FOXA1 as the top hits (p = 1×10⁻¹²), along with HNF4α and additional Forkhead family members (**Figure 3E**). These chromatin profiling results indicate that ARID1B is required to maintain active promoter- and enhancer-associated regulatory elements during sorafenib treatment, including loci enriched for hepatocyte lineage-associated transcription factor motifs.

### ARID1B loss alters transcriptional programs associated with mesenchymal transition and proliferation control

We next asked how ARID1B-dependent chromatin changes were reflected at the transcriptional level by performing RNA-seq in wild-type and ARID1B-knockout HepG2 cells treated with DMSO or 4 uM sorafenib for 72 hours (n = 3). Differential expression was modeled using DESeq2 with a genotype-by-treatment interaction design to identify transcriptional changes selectively associated with ARID1B loss during sorafenib treatment. Under sorafenib treatment, ARID1B-knockout cells exhibited 248 upregulated and 268 downregulated genes relative to wild-type controls (adjusted P < 0.1, **Figure 4A**).

**Figure 4:**
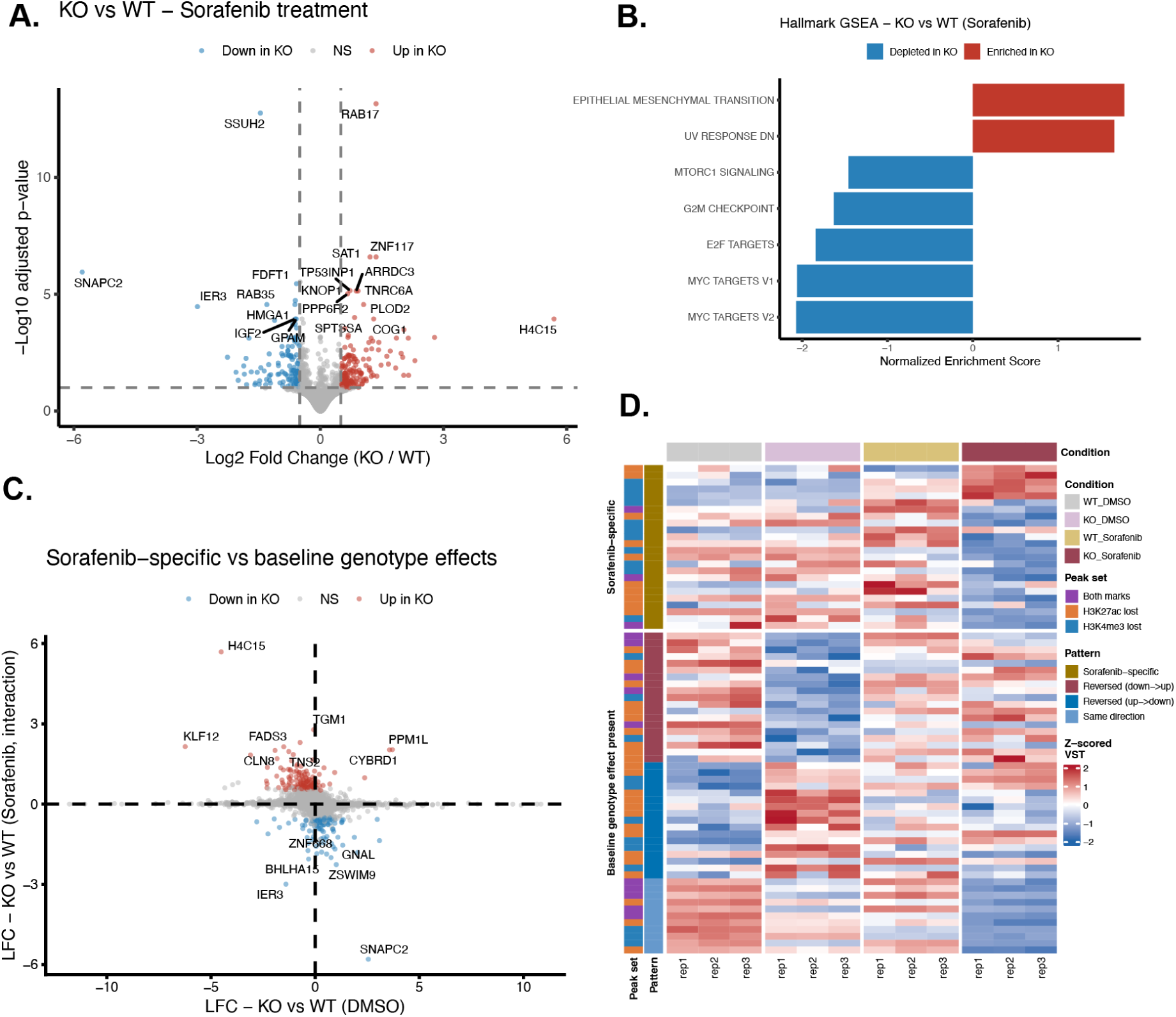
ARID1B loss induces EMT-associated transcriptional reprogramming linked to chromatin changes. **(A)** Volcano plot of RNA-seq differential expression comparing ARID1B-knockout and wild-type HepG2 cells under sorafenib treatment using the genotype-by-treatment interaction model. Significantly upregulated genes (red, n = 248) and downregulated genes (blue, n = 268) genes are highlighted (adjusted p < 0.1); non-significant genes are shown in grey. **(B)** Hallmark gene set overrepresentation analysis of ARID1B-knockout versus wild-type HepG2 cells under sorafenib treatment. Positive normalized enrichment scores indicate gene sets enriched in ARID1B-knockout cells, while negative normalized enrichment scores indicate gene sets depleted in ARID1B-knockout cells. **(C)** Scatter plot comparing log₂ fold changes from the baseline genotype effect in DMSO and the sorafenib-specific ARID1B interaction effect. Significant interaction genes are colored red if upregulated or blue if downregulated in ARID1B-knockout cells under sorafenib treatment. Selected genes are labeled. **(D)** Heatmap of VST-normalized expression for 71 significantly differentially expressed genes located within 50 kb of H3K4me3- or H3K27ac-lost peaks. Rows represent genes and columns represent biological replicates from wild-type DMSO, ARID1B-knockout DMSO, wild-type sorafenib, and ARID1B-knockout sorafenib conditions. Expression values are z-scored by gene. Row annotations indicate associated lost chromatin mark and expression pattern.

To identify pathways selectively altered in sorafenib-treated ARID1B-knockout cells, we performed gene set analysis using the MSigDB Hallmark collection. Upregulated interaction genes were enriched for epithelial-mesenchymal transition (EMT) programs, while downregulated genes were enriched for proliferative and growth-associated pathways, including MYC Targets, E2F Targets, G2M Checkpoint and mTORC1 signaling (**Figure 4B**). These changes included upregulation of EMT-associated genes such as PLOD2 and SLC2A1, and downregulating of genes including DKK1, IGF2 and HK2. Comparison of the sorafenib interaction effect with the baseline genotype effect in DMSO showed that many of these transcriptional changes were specific to the combined ARID1B loss plus sorafenib condition rather than reflecting constitutive genotype differences alone (**Figure 4C**). Overall, these results indicate that ARID1B loss rewires the transcriptional response to sorafenib, shifting cells toward a mesenchymal-like state while reducing proliferative gene programs.

To connect these transcriptional changes with the chromatin alterations identified by CUT&RUN, we linked H3K4me3- and H3K27ac-lost peaks to genes within 50 kb and intersected these genes with the RNA-seq interaction results. This identified 50 differentially expressed genes located near active chromatin peaks, including 13 genes linked to regions losing both H3K4me3 and H3K27ac. These high-confidence chromatin-linked genes included PLOD2 and SLC2A1 among upregulated genes, and DKK1, IGF2, and HK2 among downregulated genes (**Supplemental Figure S3A**). Although the 71 chromatin-linked genes are not uniformly established mediators of sorafenib resistance, several converge on pathways previously associated with multikinase inhibitor response, including glycolytic remodeling^50,51^, IGF/FGF signaling^52^, Wnt/DKK1 biology^53^ and EMT/hypoxia-associated adaptation^54,55^. This suggests that ARID1B loss may reconfigure multiple resistance-associated transcriptional programs rather than acting through a single downstream effector.

Interaction coefficients can arise from distinct underlying expression patterns; we examined normalized expression across all four genotype-treatment groups for these 71 chromatin-linked genes. 24 genes showed minimal baseline genotype effect and were altered primarily under sorafenib, while 47 genes showed a baseline genotype effect in DMSO. Among the baseline-altered genes, 36 reversed the direction of the genotype effect following sorafenib treatment, including 19 genes that shifted from lower expression in ARID1B-knockout cells at baseline to higher expression under sorafenib and 17 genes that shifted from higher expression at baseline to lower expression under sorafenib (**Figure 4D**). Representative genes illustrated multiple interaction patterns, including sorafenib-specific changes (PLOD2, IGF2), same-direction genotype effects (SLC2A1, DKK1), and reversal of genotype effects between DMSO and sorafenib conditions (HK2, KLF12) (**Supplemental Figure S3B**).

From this work, we propose that ARID1B loss reconfigures the transcriptional response to sorafenib through coordinated changes in active chromatin and gene expression. H3K27ac regions were enriched for Forkhead motifs (**Figure 3E**), so we next asked whether ARID1B loss altered FOXA transcription factor occupancy at these regulatory elements.

### FOXA1 occupancy is selectively lost in ARID1B-deficient cells during sorafenib exposure

To determine whether ARID1B loss alters lineage-associated transcription factor occupancy during therapeutic stress, we performed CUT&RUN for FOXA1 and FOXA2 in HLF cells, a second liver cancer model in which ARID1B loss confers sorafenib resistance (**Figure 2D**). Wild-type and ARID1B-knockout HLF cells were treated with DMSO or 4 μM sorafenib for 72 hours. FOXA1 occupancy was selectively reduced in ARID1B-knockout cells under sorafenib treatment, with 179 sites lost and only 1 site gained relative to wild-type cells (**Figure 5A; Supplementary Figure S4A**). In contrast, sorafenib treatment alone had minimal effect on FOXA1 binding, and only 33 FOXA1 sites were lost in ARID1B-knockout cells under DMSO conditions. Thus, the strongest FOXA1 displacement occurred only when ARID1B loss was combined with sorafenib exposure, indicating that FOXA1 occupancy is particularly vulnerable to combined genetic and therapeutic stress. FOXA2 showed a similar but more attenuated pattern, with 37 sites lost under the combined perturbation (**Supplementary Figure S4B**).

**Figure 5:**
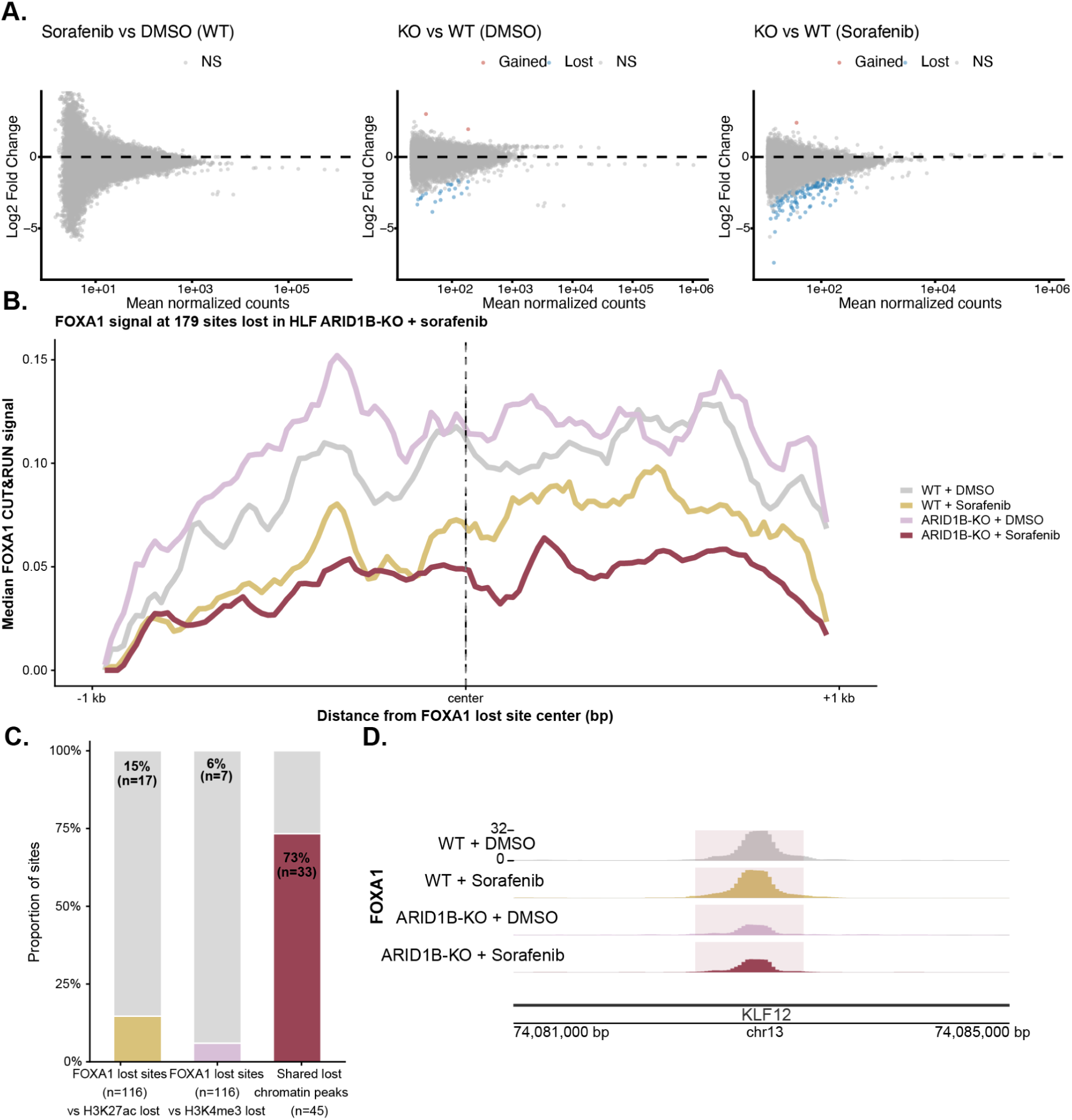
ARID1B loss selectively disrupts FOXA1 occupancy under sorafenib treatment and links FOXA loss to ARID1B-dependent active chromatin remodeling. **(A)** MA plots of FOXA1 differential occupancy in HLF cells. Left: ARID1B-KO vs. wild-type under sorafenib (179 sites lost, 1 gained). Center: ARID1B-KO vs. wild-type under DMSO (33 sites lost). Right: sorafenib vs. DMSO in wild-type cells (no significant changes). Significantly differential sites are highlighted. adjusted P < 0.1. **(B)** Median FOXA1 CUT&RUN signal at the 116 sites (adjusted P < 0.05) lost in ARID1B-KO HLF cells under sorafenib, centered on peak midpoints (±1 kb window). Lines represent the mean of two replicates per condition, smoothed with a rolling average (k = 5 bins). ARID1B-KO cells under sorafenib show selectively reduced FOXA1 occupancy relative to all other conditions. **(C)** Cross-model overlap analysis comparing FOXA-associated regulatory elements in HLF cells with ARID1B-dependent active chromatin changes in HepG2 cells. Left: overlap between shared lost H3K4me3/H3K27ac peaks from HepG2 cells and FOXA1/FOXA2 binding sites in HLF cells. Center and right: overlap between FOXA1 sites lost in ARID1B-KO HLF cells under sorafenib and H3K27ac- or H3K4me3-lost peaks from HepG2 cells. Bars show the proportion of sites in each overlap category. **(D)** Representative FOXA1 CUT&RUN signal tracks at an intronic KLF12 regulatory element (chr13:74,081,000–74,085,000, hg38) in HLF wild-type and ARID1B-KO cells treated with DMSO or sorafenib. Signal reflects scaled coverage (rep 1). FOXA1 occupancy is maintained in wild-type cells regardless of treatment but is reduced in ARID1B-KO cells under both conditions, with the greatest loss observed under sorafenib.

We next asked whether FOXA1/2 displacement in HLF cells was related to the active chromatin changes identified in HepG2 cells. Of the 45 genomic regions that lost both H3K4me3 and H3K27ac in ARID1B-knockout HepG2 cells under sorafenib treatment, 33 regions were located near FOXA1 or FOXA2 binding sites in HLF cells (73%; **Figure 5C**). This overlap links ARID1B-dependent loss of active chromatin marks to FOXA-associated regulatory elements across liver cancer models. Conversely, only a subset of FOXA1-lost sites in HLF cells overlapped HepG2 regions losing active histone marks: 17 FOXA1-lost sites overlapped H3K27ac-lost peaks and 7 overlapped H3K4me3-lost peaks. These results suggest that FOXA displacement and active histone mark depletion are related but not identical regulatory events, likely reflecting both shared ARID1B-dependent regulatory programs and cell-line-specific chromatin contexts.

To determine whether FOXA1 displacement was associated with transcriptional changes, we examined expression of genes proximal to FOXA1-lost sites. Genes near FOXA1-lost sites showed a modest but significant upward shift in expression relative to background genes (Wilcoxon p = 0.016; **Supplementary Figure S4C**), consistent with a context-dependent role for FOXA1 in both transcriptional activation and repression. Among significantly altered genes near FOXA1-lost sites, KLF12 was robustly induced (log2FC = 2.15, padj = 0.030), whereas CCN2/CTGF was repressed (log2FC = −0.57, padj = 0.044; **Supplementary Figure S4D**). These examples support the idea that FOXA1 loss does not produce a uniform transcriptional direction, but instead contributes to regulatory rewiring at distinct chromatin contexts.

We then examined FOXA1 occupancy at an intronic regulatory element within KLF12, the most significantly upregulated gene proximal to FOXA1-lost sites. FOXA1 binding at this locus was robust in wild-type cells under both DMSO and sorafenib treatment, but was reduced in ARID1B-knockout cells, with the greatest loss observed after sorafenib exposure (**Figure 5D**). Because KLF12 induction has been linked to invasive and drug-resistant transcriptional states, this locus provides a representative example of ARID1B-dependent FOXA1 destabilization at a regulatory element associated with sorafenib resistance.

These results support a model in which ARID1B helps preserve sorafenib sensitivity by maintaining active chromatin and FOXA pioneer factor occupancy at lineage-associated regulatory elements. Loss of ARID1B under therapeutic stress disrupts this regulatory architecture, promoting chromatin and transcriptional reprogramming toward EMT-associated resistance states.

## DISCUSSION

Our findings identify cBAF as a critical determinant of sorafenib sensitivity in hepatocellular carcinoma and define a chromatin-based mechanism through which cBAF loss promotes therapeutic resistance. Using an unbiased CRISPR screen targeting epigenetic regulators, we identified ARID1A, ARID1B, and SMARCC1, core components of the cBAF complex, as among the strongest resistance-associated hits. In contrast, ARID2, a subunit of the related PBAF complex, had little effect. This specificity implicates cBAF rather than SWI/SNF disruption broadly, consistent with emerging evidence that distinct SWI/SNF assemblies perform non-redundant functions in cancer.

The clinical relevance of this finding is supported by TCGA survival data showing that HCCs harboring ARID1A or ARID1B alterations have significantly worse overall survival than tumors without these alterations. Because these data are correlative and do not directly measure response to sorafenib, they should not be interpreted as evidence that ARID1A/B mutations predict sorafenib resistance in patients. Instead, the survival data raises the clinically actionable hypothesis that cBAF alteration status may identify a subset of HCCs with poor prognosis and altered therapeutic vulnerability.

Mechanistically, we show that ARID1B loss under sorafenib exposure triggers selective collapse of active chromatin at a discrete set of regulatory loci. These regions lose both H3K4me3 and H3K27ac, consistent with coordinated decommissioning of active promoters and enhancers rather than nonspecific transcriptional noise. Because ARID1B is a defining subunit of the cBAF complex, we interpret these sites as cBAF-dependent regulatory elements whose active chromatin state requires intact cBAF activity under drug-induced stress. This model is directly supported by our finding that 53 of 74 shared lost peaks, or 72%, are occupied by ARID1B in wild-type cells and lose ARID1B occupancy in knockout cells. These results are consistent with displacement of cBAF from ARID1B-dependent regulatory loci rather than purely indirect chromatin effects. The greater magnitude of H3K27ac loss compared with H3K4me3 further suggests that enhancer elements may be particularly dependent on cBAF for maintenance during sorafenib exposure, consistent with the established role of SWI/SNF complexes in preserving nucleosome-depleted regulatory regions^9^.

Motif analysis of H3K27ac-lost peaks revealed strong enrichment for FOXA family motifs, with FOXA1 emerging as the top individual motif, alongside HNF4α, a master regulator of hepatocyte identity. Both factors are pioneer transcription factors that establish and maintain hepatocyte-specific regulatory landscapes. Their enrichment at cBAF-dependent lost sites suggested that ARID1B loss may destabilize hepatocyte pioneer factor-associated regulatory elements under drug stress. Consistent with this hypothesis, CUT&RUN profiling in HLF cells identified 179 FOXA1 sites that were lost specifically in ARID1B-KO cells under sorafenib treatment, with limited effects of either perturbation alone. The synthetic nature of this effect, requiring both ARID1B loss and sorafenib exposure, supports a model in which cBAF becomes particularly important for maintaining FOXA1-associated regulatory architecture under kinase inhibitor stress.

The transcriptional consequences of these chromatin changes were consistent with a shift away from a differentiated, proliferative hepatocyte-like state. Genes linked to FOXA1-lost sites showed a modest but statistically significant increase in expression compared with background, consistent with the context-dependent ability of FOXA1 to function as either a transcriptional activator or repressor. Among individual linked genes, KLF12 was robustly induced and appeared in both the shared chromatin-loss and FOXA1-loss analyses. KLF12 is a Krüppel-like factor with reported roles in epithelial–mesenchymal transition and invasion, suggesting that its induction may contribute to the mesenchymal transcriptional program observed by gene set enrichment analysis. In contrast, DKK1, a Wnt pathway inhibitor, and CCN2/CTGF, a TGF-β-responsive gene, were repressed. These observations nominate a small yet mechanistically coherent set of downstream effectors that may connect FOXA1 displacement to transcriptional reprogramming during sorafenib resistance.

Transcriptome-wide, ARID1B loss under sorafenib treatment was associated with induction of epithelial–mesenchymal transition programs and suppression of MYC targets, E2F targets, G2M checkpoint genes, and mTORC1 signaling. This pattern is consistent with a transition from an actively proliferative, drug-sensitive state toward a more quiescent, mesenchymal-like drug-tolerant state. Such persister-like states are increasingly recognized as mechanisms of resistance to targeted therapies across cancer types^56^. Because sorafenib targets RAF–MEK–ERK and VEGFR signaling,^21,23,24,50–53^ suppression of proliferative transcriptional programs may reflect adaptive rewiring in response to kinase inhibition rather than simple growth arrest^57^.

This work supports models in which cBAF complex integrity is required to sustain the hepatocyte pioneer factor network during kinase inhibitor stress^5^. When cBAF function is lost, sorafenib exposure reveals a chromatin maintenance vulnerability at FOXA1/HNF4α-associated regulatory elements, triggering epigenetic reprogramming toward a mesenchymal-like resistance state. These findings nominate the cBAF–FOXA1 regulatory axis as a candidate mechanism of sorafenib resistance in HCC and suggest that strategies aimed at preserving hepatocyte regulatory identity or targeting EMT-associated adaptive states may improve therapeutic responses in cBAF-deficient tumors.

## Acknowledgements

TCGA data were obtained through cBioPortal. The Graphical Abstract, as well as Figures 1A and 3A were created in <u>BioRender.com</u>. Claude AI was used to compile Supplemental Data into a spreadsheet.

## Author contributions

Jacqueline A Brinkman: Formal analysis, Investigation, Data curation, Visualization, Methodology, Software, Validation, Writing—original draft. Margarita Dzama-Karels, Ph.D.: Investigation, Formal analysis, Methodology, Visualization, Validation, Writing—review & editing. Max Bucklan: Data curation, Validation; Carl Manner: Investigation, Data curation; Mallory Sokolowski: Visualization; Jesse Raab, Ph.D.: Conceptualization, Investigation, Formal analysis, Data curation, Methodology, Visualization, Software, Supervision, Writing—review & editing.

## Conflict of interest

The authors declare no conflicts of interest.

## Funding

This work was supported by the Lineberger Comprehensive Cancer Center to Jesse Raab, Ph.D National Institute of General Medical Sciences of the National Institutes of Health to Jacqueline A Brinkman under Award Number T32 [1T32GM135128]; the American Cancer Society to Margarita Dzama-Karels, Ph.D [PF-24-1247688-01-DMC]; and National Institute of General Medical Sciences of the National Institutes of Health to Mallory Sokolowski under Award Number T32 [T32GM135128].

## Data availability

All high-throughput sequencing data will be available in Gene Expression Omnibus (GEO) upon acceptance:

---

**Supplemental Figure S1:**
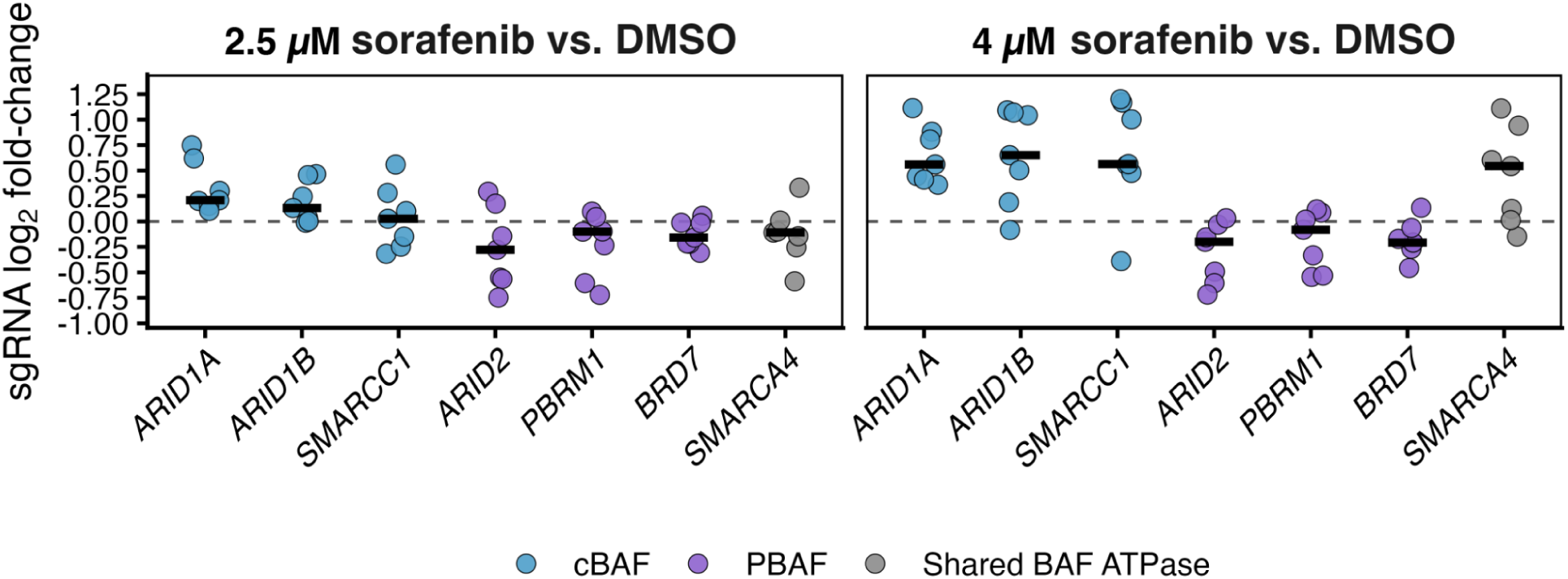
cBAF, but not PBAF, subunit sgRNAs are enriched under sorafenib selection. Individual sgRNA-level log₂ fold-change for selected BAF complex subunits in 2.5 uM and 4 uM sorafenib versus DMSO comparisons. Each point represents one sgRNA targeting the indicated gene, and crossbars indicate the median sgRNA log₂ fold-change for each gene. sgRNAs targeting cBAF subunits (ARID1A, ARID1B and SMARCC1; blue) were enriched under sorafenib selection, whereas sgRNAs targeting the PBAF-specific subunits ARID2, PBRM1 and BRD7 (purple) were not. The shared BAF ATPase SMARCA4 is shown in grey. Positive values indicate sgRNA enrichment under sorafenib selection.

**Supplemental Figure S2:**
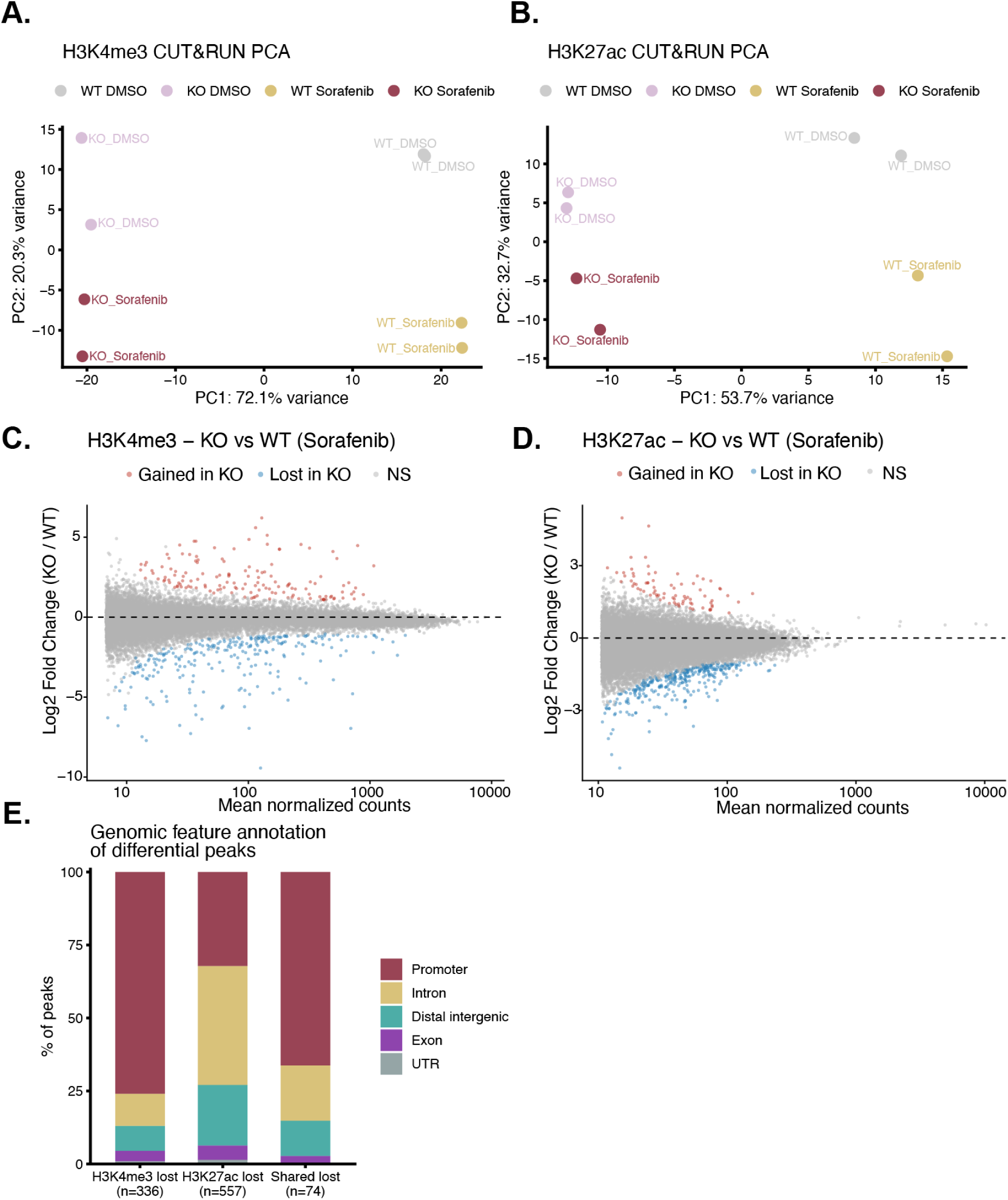
ARID1B loss remodels active chromatin landscapes during sorafenib treatment. **(A)** Principal component analysis of H3K4me3 CUT&RUN samples. **(B)** Principal component analysis of H3K27ac CUT&RUN samples. **(C)** MA plot of H3K4me3 differential peak analysis comparing ARID1B-knockout and wild-type HepG2 cells under sorafenib treatment. Significantly gained peaks (red, n = 180) and lost peaks (blue, n = 336) are highlighted; non-significant peaks are shown in grey. **(D)** MA plot of H3K27ac differential peak analysis comparing ARID1B-knockout and wild-type HepG2 cells under sorafenib treatment. Significantly gained peaks (red, n = 124) and lost peaks (blue, n = 557) are highlighted; non-significant peaks are shown in grey. **(E)** Genomic feature annotation of H3K4me3 lost, H3K27ac lost and shared lost peak sets. H3K4me3 lost peaks are predominantly promoter-associated and H3K27ac lost peaks are enriched at intronic and distal intergenic regions.

**Supplemental Figure S3:**
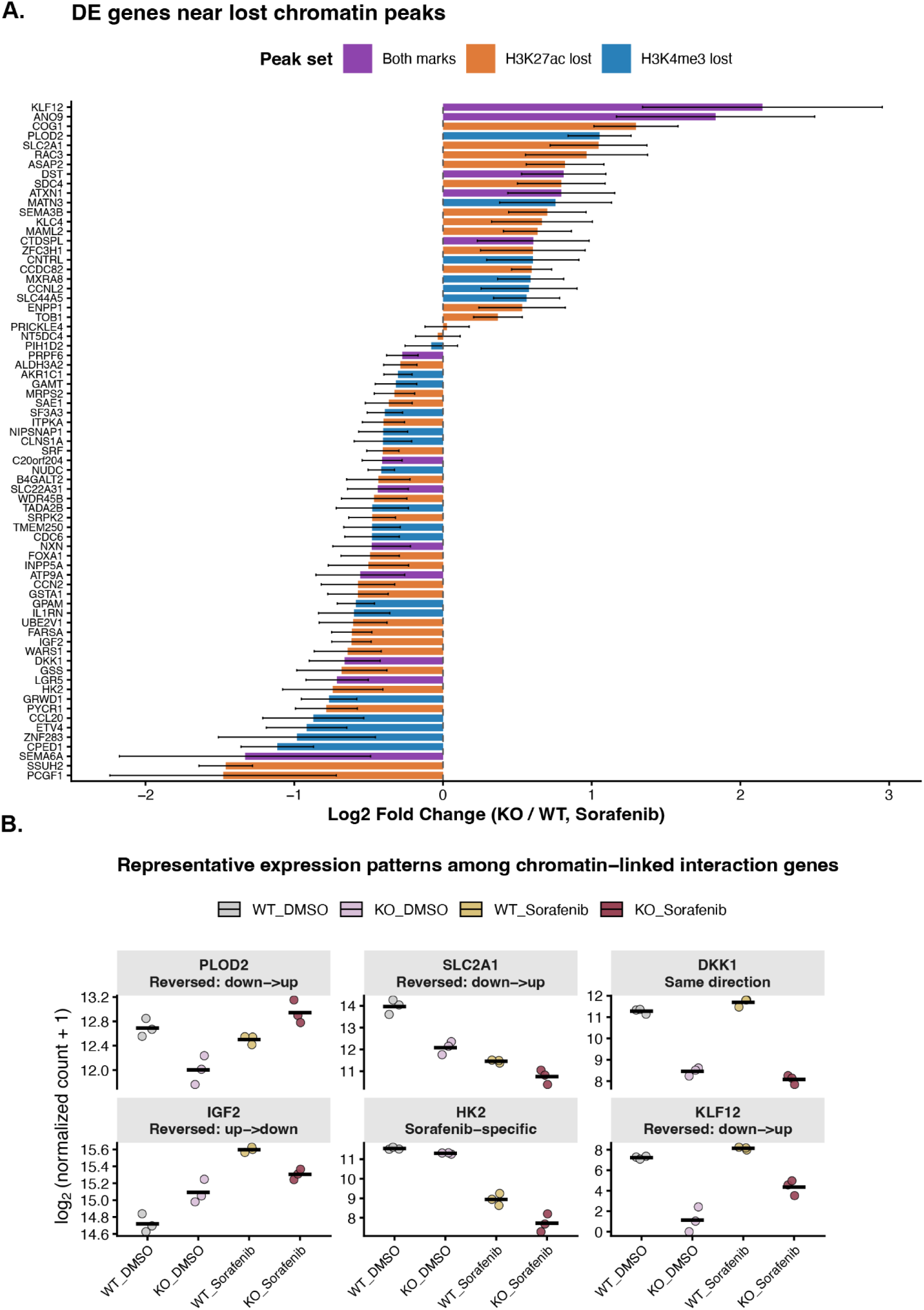
Chromatin-linked ARID1B-sorafenib interaction genes show distinct baseline and drug-dependent expression patterns. (**A)** log₂ fold changes for significantly differentially expressed genes located within 100 kb of H3K4me3- or H3K27ac-lost peaks. Bars are colored by the associated peak set: purple, genes linked to regions losing both marks; orange, H3K27ac-lost peaks only; blue, genes linked to H3K4me3-lost peaks only. Error bars represent ±1 standard error of the shrunken log₂ fold-change estimate. **(B)** Representative chromatin-linked interaction genes plotted as log₂-transformed DESeq2-normalized counts across wild-type DMSO, ARID1B-knockout DMSO, wild-type sorafenib, and ARID1B-knockout sorafenib conditions. Points represent biological replicates, and horizontal bars indicate group means. Genes were selected to illustrate major expression patterns observed among chromatin-linked interaction genes, including sorafenib-specific effects, same-direction genotype effects, and reversal of genotype effects between DMSO and sorafenib.

**Supplemental Figure S4:**
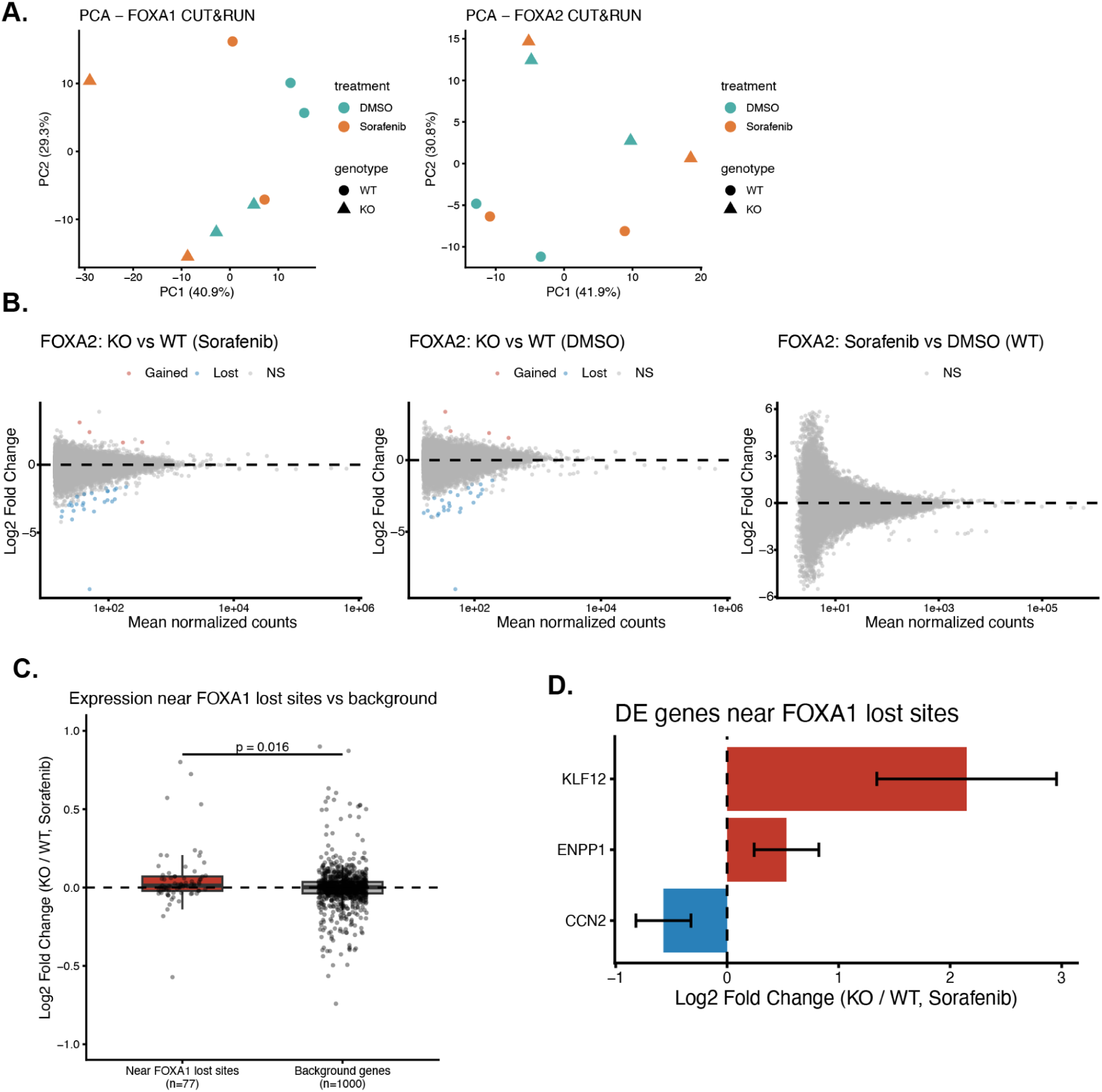
FOXA2 occupancy changes and transcriptional associations at FOXA1 lost sites. (**A)** Principal component analysis of FOXA1 (left) and FOXA2 (right) CUT&RUN signal from HLF wild-type and ARID1B-KO cells treated with DMSO or sorafenib (n = 2 replicates per condition). **(B)** MA plots of FOXA2 differential occupancy. Left: ARID1B-KO vs. wild-type under sorafenib (37 sites lost, 5 gained). Center: ARID1B-KO vs. wild-type under DMSO. Right: sorafenib vs. DMSO in wild-type cells. **(C)** Boxplot of log2 fold changes (KO vs. WT, sorafenib) for genes proximal to FOXA1 lost sites (n = 77) compared to a random background gene set (n = 1,000). Two-sided Wilcoxon rank-sum test, p = 0.016. **(D)** Log2 fold changes for significantly differentially expressed genes proximal to FOXA1 lost sites. Error bars indicate ±1 SE of the shrunken log2 fold change estimate. KLF12 (log2FC = 2.15, padj = 0.030) and CCN2 (log2FC = −0.57, padj = 0.044) are highlighted as the most significant induced and repressed genes, respectively.

## Notes

### Competing Interest Statement

The authors have declared no competing interest.

## References

1. Kadoch, C. et al. Proteomic and bioinformatic analysis of mammalian SWI/SNF complexes identifies extensive roles in human malignancy. Nat Genet 45, 592–601 (2013).

2. Shain, A. H. & Pollack, J. R. The spectrum of SWI/SNF mutations, ubiquitous in human cancers. PLoS One 8, e55119 (2013).

3. Michel, B. C. et al. A non-canonical SWI/SNF complex is a synthetic lethal target in cancers driven by BAF complex perturbation. Nat Cell Biol 20, 1410–1420 (2018).

4. Ho, L. & Crabtree, G. R. Chromatin remodelling during development. Nature 463, 474–484 (2010).

5. Aoki, K. et al. Canonical BAF complex regulates the oncogenic program in human T-cell acute lymphoblastic leukemia. Blood 143, 604–618 (2024).

6. Nocente, M. C. et al. cBAF generates subnucleosomes that expand OCT4 binding and function beyond DNA motifs at enhancers. Nat Struct Mol Biol 31, 1756–1768 (2024).

7. Soleil, M. et al. How cBAF and PBAF regulate the nucleosomal and subnucleosomal landscape of promoters, enhancers and REST repressor binding sites. 2025.06.12.659349 Preprint at 10.1101/2025.06.12.659349 (2025).

8. Mashtalir, N. et al. Modular Organization and Assembly of SWI/SNF Family Chromatin Remodeling Complexes. Cell 175, 1272–1288.e20 (2018).

9. Wang, W. et al. Diversity and specialization of mammalian SWI/SNF complexes. Genes Dev 10, 2117–2130 (1996).

10. Versteege, I. et al. Truncating mutations of hSNF5/INI1 in aggressive paediatric cancer. Nature 394, 203–206 (1998).

11. Wiegand, K. C. et al. ARID1A mutations in endometriosis-associated ovarian carcinomas. N Engl J Med 363, 1532–1543 (2010).

12. Roberts, C. W. M., Leroux, M. M., Fleming, M. D. & Orkin, S. H. Highly penetrant, rapid tumorigenesis through conditional inversion of the tumor suppressor gene Snf5. Cancer Cell 2, 415–425 (2002).

13. Davoli, T. et al. Cumulative haploinsufficiency and triplosensitivity drive aneuploidy patterns and shape the cancer genome. Cell 155, 948–962 (2013).

14. Helming, K. C., Wang, X. & Roberts, C. W. M. Vulnerabilities of mutant SWI/SNF complexes in cancer. Cancer Cell 26, 309–317 (2014).

15. Mittal, P. & Roberts, C. W. M. The SWI/SNF complex in cancer - biology, biomarkers and therapy. Nat Rev Clin Oncol 17, 435–448 (2020).

16. Jing, H., Zhang, X. & Meng, L. Swinging the SWI/SNF complexes for cancer therapy. Trends in Pharmacological Sciences 46, 907–921 (2025).

17. Sung, H. et al. Global Cancer Statistics 2020: GLOBOCAN Estimates of Incidence and Mortality Worldwide for 36 Cancers in 185 Countries. CA Cancer J Clin 71, 209–249 (2021).

18. Llovet, J. M. et al. Sorafenib in advanced hepatocellular carcinoma. N Engl J Med 359, 378–390 (2008).

19. Tang, W. et al. The mechanisms of sorafenib resistance in hepatocellular carcinoma: theoretical basis and therapeutic aspects. Sig Transduct Target Ther 5, 87 (2020).

20. Wei, L. et al. Genome-wide CRISPR/Cas9 library screening identified PHGDH as a critical driver for Sorafenib resistance in HCC. Nat Commun 10, 4681 (2019).

21. Tang, W. et al. The mechanisms of sorafenib resistance in hepatocellular carcinoma: theoretical basis and therapeutic aspects. Sig Transduct Target Ther 5, 87 (2020).

22. Zhou, Y. et al. GPAT3 is a potential therapeutic target to overcome sorafenib resistance in hepatocellular carcinoma. Theranostics 14, 3470–3485 (2024).

23. Gao, R. et al. YAP/TAZ and ATF4 drive resistance to Sorafenib in hepatocellular carcinoma by preventing ferroptosis. EMBO Mol Med 13, e14351 (2021).

24. Zheng, A. et al. CRISPR/Cas9 genome-wide screening identifies KEAP1 as a sorafenib, lenvatinib, and regorafenib sensitivity gene in hepatocellular carcinoma. Oncotarget 10, 7058–7070 (2019).

25. Rudalska, R. et al. In vivo RNAi screening identifies a mechanism of sorafenib resistance in liver cancer. Nat Med 20, 1138–1146 (2014).

26. Preziosi, M. E. et al. In vivo screen identifies LXR agonism potentiates sorafenib killing of hepatocellular carcinoma. 668350 Preprint at 10.1101/668350 (2019).

27. Cancer Genome Atlas Research Network. Electronic address: & Cancer Genome Atlas Research Network. Comprehensive and Integrative Genomic Characterization of Hepatocellular Carcinoma. Cell 169, 1327–1341.e23 (2017).

28. Schulze, K. et al. Exome sequencing of hepatocellular carcinomas identifies new mutational signatures and potential therapeutic targets. Nat Genet 47, 505–511 (2015).

29. Iwafuchi-Doi, M. & Zaret, K. S. Cell fate control by pioneer transcription factors. Development 143, 1833–1837 (2016).

30. Zaret, K. S. & Carroll, J. S. Pioneer transcription factors: establishing competence for gene expression. Genes Dev 25, 2227–2241 (2011).

31. Mathur, R. et al. ARID1A loss impairs enhancer-mediated gene regulation and drives colon cancer in mice. Nat Genet 49, 296–302 (2017).

32. Alver, B. H. et al. The SWI/SNF chromatin remodelling complex is required for maintenance of lineage specific enhancers. Nat Commun 8, 14648 (2017).

33. Joung, J. et al. Genome-scale CRISPR-Cas9 knockout and transcriptional activation screening. Nat Protoc 12, 828–863 (2017).

34. Li, W. et al. MAGeCK enables robust identification of essential genes from genome-scale CRISPR/Cas9 knockout screens. Genome Biol 15, 554 (2014).

35. Patro, R., Duggal, G., Love, M. I., Irizarry, R. A. & Kingsford, C. Salmon provides fast and bias-aware quantification of transcript expression. Nature Methods 14, 417–419 (2017).

36. Love, M. I. et al. Tximeta: Reference sequence checksums for provenance identification in RNA-seq. PLOS Computational Biology 16, e1007664 (2020).

37. Love, M. I., Huber, W. & Anders, S. Moderated estimation of fold change and dispersion for RNA-seq data with DESeq2. Genome Biology 15, 550 (2014).

38. Zhu, A., Ibrahim, J. G. & Love, M. I. Heavy-tailed prior distributions for sequence count data: removing the noise and preserving large differences. Bioinformatics 35, 2084–2092 (2019).

39. Korotkevich, G. et al. Fast gene set enrichment analysis. (2016) doi:10.1101/060012.

40. Meers, M. P., Bryson, T. D., Henikoff, J. G. & Henikoff, S. Improved CUT&RUN chromatin profiling tools. eLife 8, e46314 (2019).

41. Dzama-Karels, M. et al. Menin-MLL1 complex cooperates with NF-Y to promote hepatocellular carcinoma survival. Cell Rep 44, 116619 (2025).

42. Brinkman, J. A., Hantelys, F., Raab, J. & Gracz, A. D. Chromatin State Distinguishes Injury-Responsive from State-Stabilizing Transcriptional Programs in Hybrid Hepatocytes. 2026.05.04.722673 Preprint at 10.64898/2026.05.04.722673 (2026).

43. Langmead, B. & Salzberg, S. L. Fast gapped-read alignment with Bowtie 2. Nat Methods 9, 357–359 (2012).

44. Jalili, V., Matteucci, M., Masseroli, M. & Morelli, M. J. Using combined evidence from replicates to evaluate ChIP-seq peaks. Bioinformatics 31, 2761–2769 (2015).

45. Lun, A. T. L. & Smyth, G. K. csaw: a Bioconductor package for differential binding analysis of ChIP-seq data using sliding windows. Nucleic Acids Res 44, e45 (2016).

46. Ramírez, F. et al. deepTools2: a next generation web server for deep-sequencing data analysis. Nucleic Acids Res 44, W160–W165 (2016).

47. ggplot2: Elegant Graphics for Data Analysis (3e). https://ggplot2-book.org/.

48. Slowikowski K (2026). ggrepel: Automatically Position Non-Overlapping Text Labels with ‘ggplot2’. R package version 0.9.8,.

49. Kramer, N. E. et al. Plotgardener: cultivating precise multi-panel figures in R. Bioinformatics 38, 2042–2045 (2022).

50. Leoni, I. et al. MiR-22/GLUT1 Axis Induces Metabolic Reprogramming and Sorafenib Resistance in Hepatocellular Carcinoma. Int J Mol Sci 26, 3808 (2025).

51. Tesori, V. et al. The multikinase inhibitor Sorafenib enhances glycolysis and synergizes with glycolysis blockade for cancer cell killing. Sci Rep 5, 9149 (2015).

52. Tovar, V. et al. Tumour initiating cells and IGF/FGF signalling contribute to sorafenib resistance in hepatocellular carcinoma. Gut 66, 530–540 (2017).

53. Seo, S. H. et al. Inhibition of Dickkopf-1 enhances the anti-tumor efficacy of sorafenib via inhibition of the PI3K/Akt and Wnt/β-catenin pathways in hepatocellular carcinoma. Cell Commun Signal 21, 339 (2023).

54. Cai, H., Hu, G., Chen, J., Feng, H. & Bi, Y. The role of PLOD family genes in liver hepatocellular carcinoma: from mechanisms to therapeutic potential. BMC Cancer 26, 172 (2026).

55. Noda, T. et al. PLOD2 induced under hypoxia is a novel prognostic factor for hepatocellular carcinoma after curative resection. Liver Int 32, 110–118 (2012).

56. Wang, Z. et al. Drug-tolerant persister cells in cancer: bridging the gaps between bench and bedside. Nat Commun 16, 10048 (2025).

57. East, M. P. & Johnson, G. L. Adaptive chromatin remodeling and transcriptional changes of the functional kinome in tumor cells in response to targeted kinase inhibition. J Biol Chem 298, 101525 (2021).

